## Supplementary Figures for "HIFα isoform specific activities drive cell-type specificity of *VHL*-associated oncogenesis"

### Supplementary Fig. 1

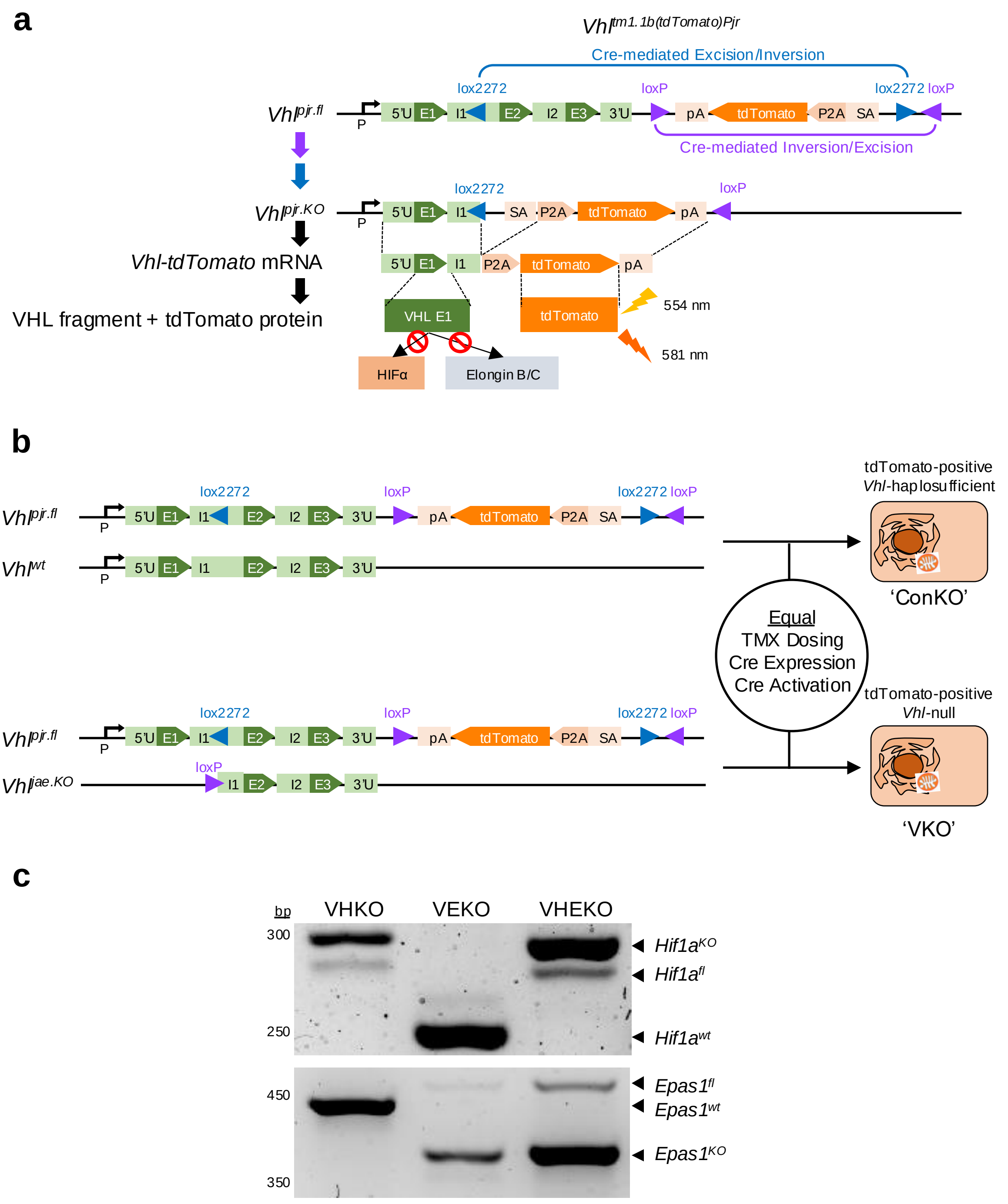


**Supplementary Fig. 1: Co-deletion of *Hif1a* and *Epas1* in tdTomato-tagged *Vhl*-recombined cells. (a)** Schematic diagram depicting the design and recombination of the *Vhl*$^{pjr.fl}$ allele. *Vhl*$^{pjr.fl}$ and *Vhl*$^{pjr.KO}$ refer to 'floxed' and 'knockout' forms of the *Vhl*$^{pjr}$ allele. P, *Vhl* promoter; U, untranslated region; E, *Vhl* exon; I, *Vhl* intron; pA, polyadenylation site; P2A, porcine teschovirus 2A peptide; SA, splice acceptor. Dashed lines - spliced and translated regions; lightning symbols - excitation and emission wavelengths for tdTomato fluorescence. Red crossed circle indicates no interaction between VHL exon 1 fragment and HIFα or Elongin B/C. **(b)** *Vhl* allele construction for ConKO and VKO mice. Mice of both genotypes carry the *Vhl*$^{pjr.fl}$ allele. ConKO mice carry a second wild-type *Vhl*$^{wt}$ allele, while VKO mice carry a constitutively inactivated *Vhl*$^{jae.KO}$ allele. **(c)** Genomic PCR for *Hif1a* and *Epas1* alleles performed on FAC-sorted tdTomato-positive cells from VHKO, VEKO, and VHEKO mice given 5x 2 mg tamoxifen and harvested early (1-3 weeks) after recombination. Expected amplicon sizes for wild-type (wt), floxed/unrecombined (fl) and knock-out/recombined (KO) forms of *Hif1a* and *Epas1* depicted by black arrows.

### Supplementary Fig. 2

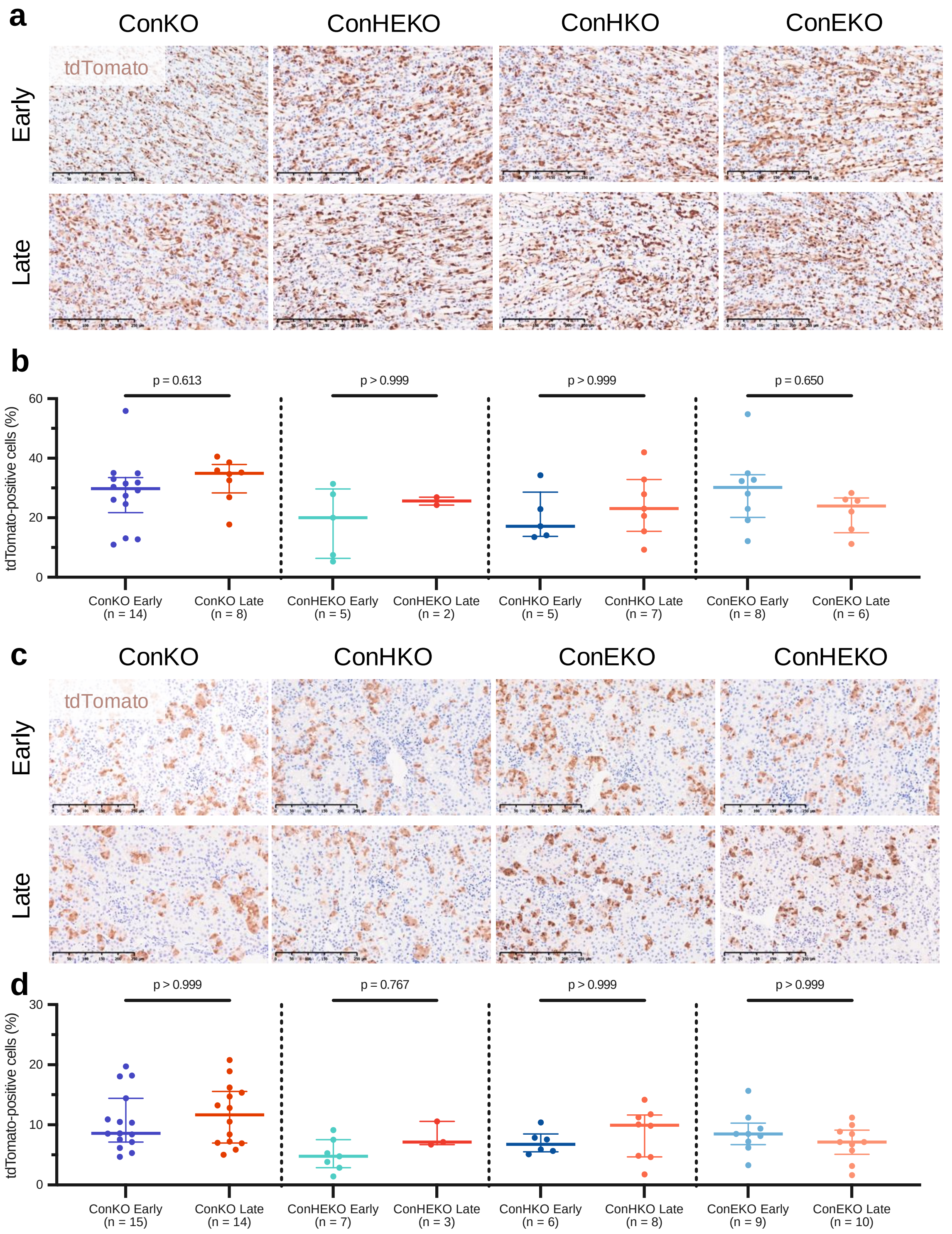


**Supplementary Fig. 2: Response to HIFα inactivation in different renal regions of 'control' kidneys. (a, c)** Representative tdTomato IHC counterstained with hematoxylin in the renal papilla **(a)** or renal cortex and outer medulla **(c)** of ConKO, ConHKO, ConEKO, and ConHEKO mice given 5x 2 mg tamoxifen and harvested early (1-3 weeks) or late (4-12 months) following recombination. Scale bar denotes 250 µm; 20x magnification. **(b, d)** Automated quantification (see **Methods**) of the proportion of cells that are tdTomato-positive in the renal papilla **(b)** or renal cortex and outer medulla **(d).** Pairwise comparisons by Kruskal-Wallis test with Dunn's correction. Median and inter-quartile range plotted.

#### Supplementary Fig. 3

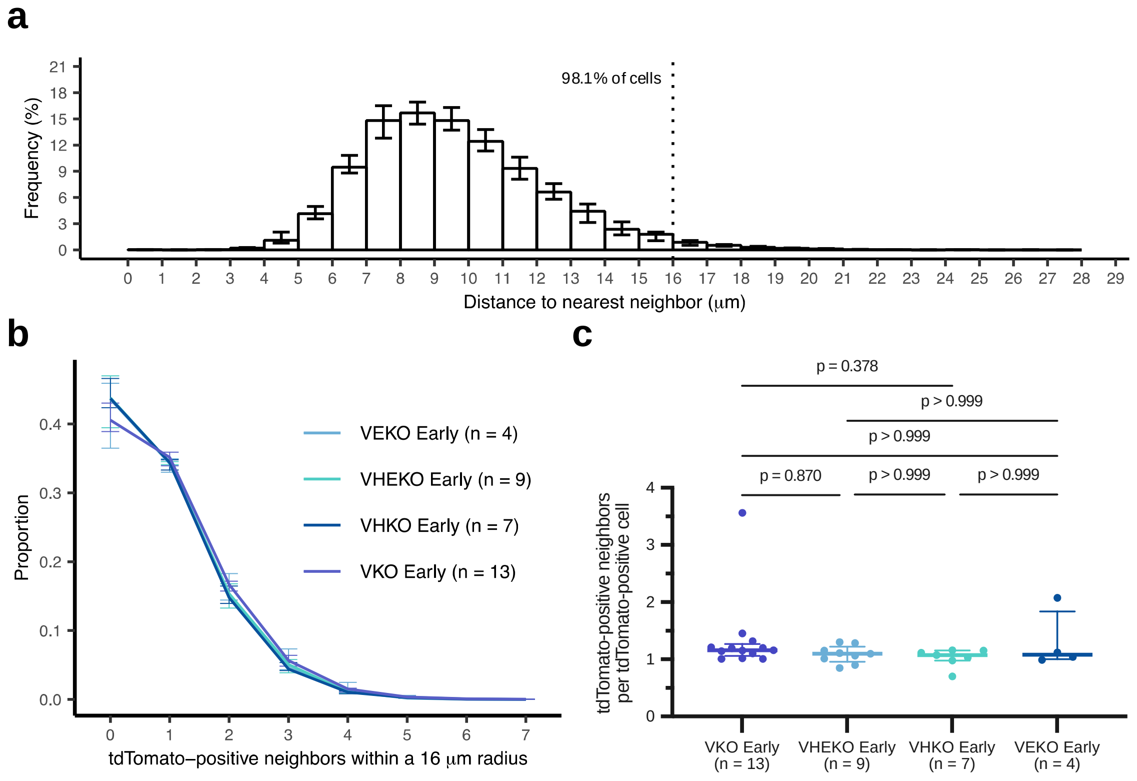


**Supplementary Fig. 3: Spatial distribution of cells in the renal cortex and outer medulla. (a)** Histogram depicting the distance to the nearest neighbor, whether tagged or untagged, for tdTomato-positive cells in the renal cortex and outer medulla. Median and interquartile range plotted. 50,000 cells analyzed over n = 10 mice chosen randomly across all genotypes and timepoints. **(b, c)** Frequency distribution **(b)**, and mean number **(c)** of tdTomato-positive neighbors of tdTomato-positive cells within a 16 μm radius in the cortex and outer medulla of VKO, VHEKO, VHKO, and VEKO mice harvested early after *Vhl* inactivation. Median and inter-quartile range plotted. Pairwise comparisons by Kruskal-Wallis test with Dunn's correction.

#### Supplementary Fig. 4

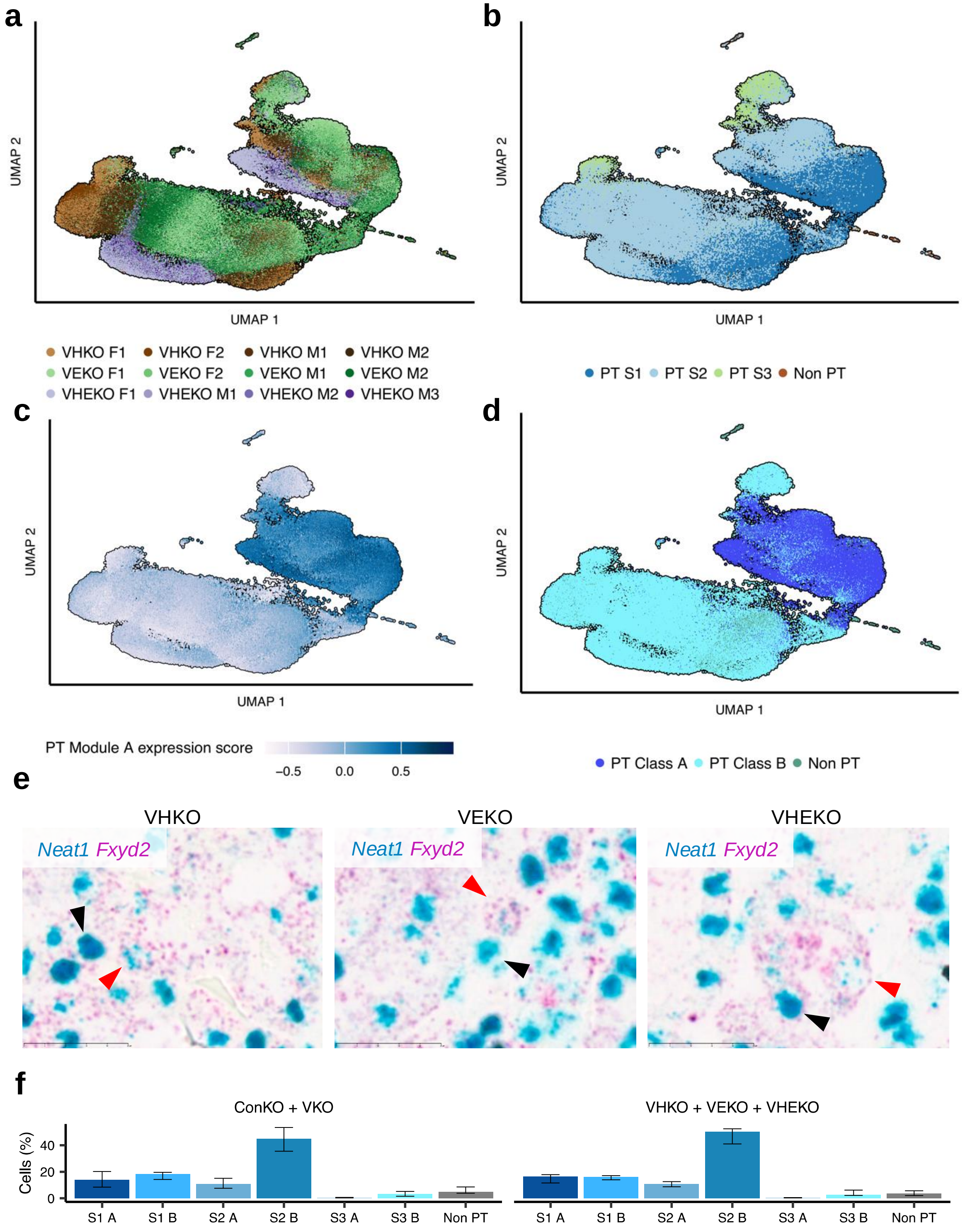


**Supplementary Fig. 4: Single-cell RNA sequencing of tdTomato-tagged cells from VHKO, VEKO, and VHEKO mice. (a-d)** UMAP plots depicting tdTomato-positive cells of n = 4 VHKO, VEKO, and VHEKO mice harvested at 4-12 months following *Vhl* and *Hif1a* and/or *Epas1* inactivation. **(a)** Data distribution by individual mouse. Cell colors are shaded according to the individual mouse within the genotypic group (VHKO in shades of brown; VEKO in shades of green; VHEKO in shades of purple). Male (M); Female (F). Concordance of the shades of the colors in UMAP space illustrates the reproducibility of the data. **(b)** Data distribution by assigned renal cell type. Proximal Tubule (PT). Cells of similar cell types are positioned together in UMAP space. **(c)** Expression score for PT Module A. High and low expression of PT Module A associates with distinct areas in UMAP space. **(d)** Data distribution by PT class as defined by PT Module A expression score. **(e)** Representative dual RNA *in situ* hybridization for *Neat1* (blue) and *Fxyd2* (pink) mRNA in kidney sections from VHKO, VEKO, and VHEKO mice. PT Class A (*Neat1*$^{high}$/*Fxyd2*$^{low}$; black arrows) and PT Class B (*Neat1*$^{low}$/*Fxyd2*$^{high}$; red arrows) cells are indicated. **(f)** Proportion of cells (median and inter-quartile range) that are of each PT identity in scRNA-seq datasets of ConKO and VKO (left), or VHKO, VEKO, and VHEKO (right) cells. Both datasets exhibit similar composition of PT identities. **(b, d)** All cell types other than PT S1, S2, and S3 cell types are assigned as 'Non PT'.

#### Supplementary Fig. 5

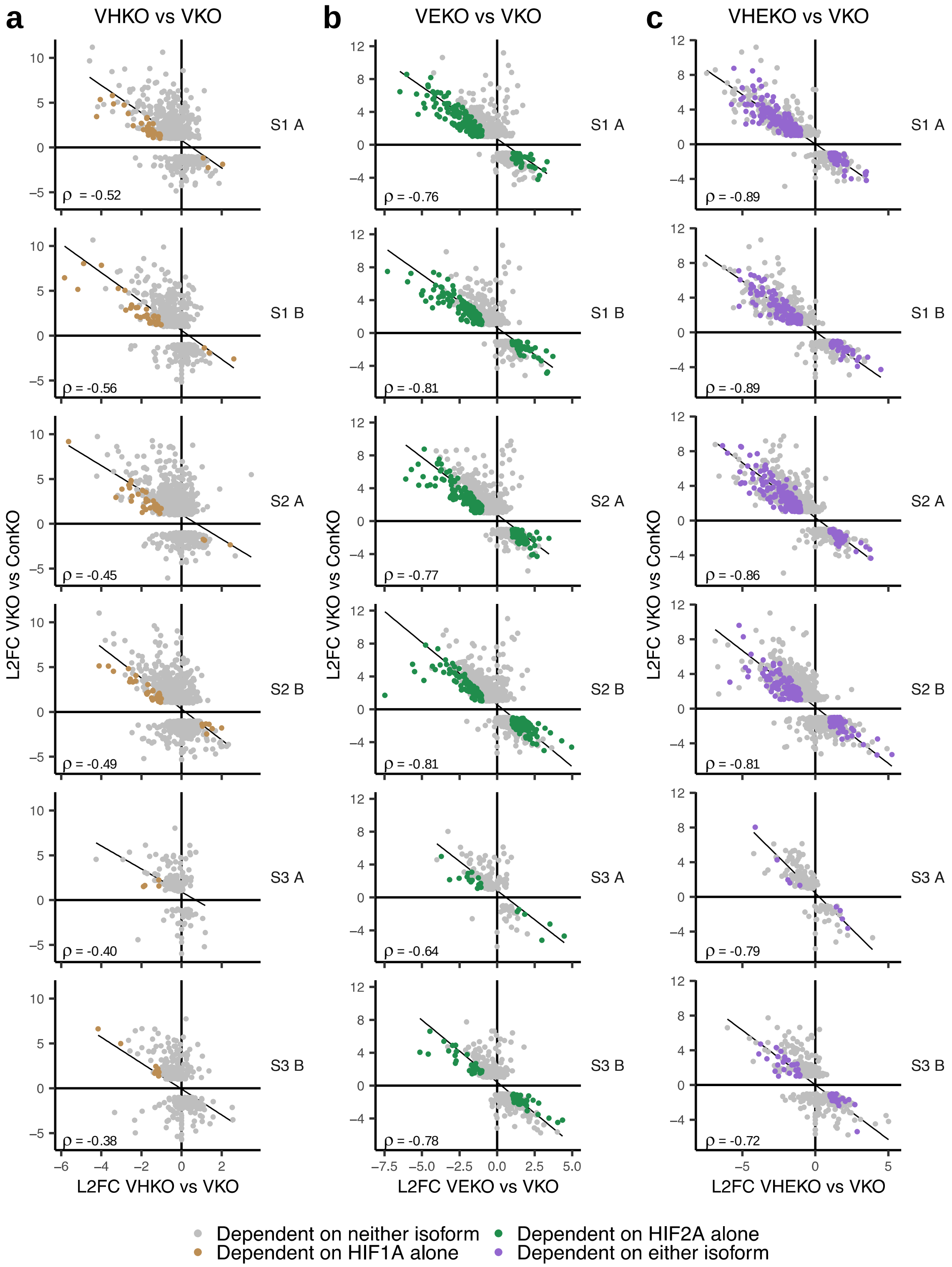


**Supplementary Fig. 5: HIFα isoform-specific dependence of gene expression in *Vhl*-null cells. (a-c)** Scatter plots depicting the effects of individual or combined HIFα co-deletion on expression of *Vhl*-dependent genes in each PT identity. Each scatter plot depicts pseudo-bulked $\log_2$-fold changes (L2FC) for *Vhl*-dependent genes in the indicated VHKO **(a)**, VEKO **(b)**, and VHEKO **(c)** combined *Vhl*/HIFα-null mice *versus* VKO mice. Spearman's correlation coefficients (ρ) for each comparison are indicated. Genes are colored by the HIFα-dependence of their regulation. Genes whose regulation was reversed only after combined HIF deletion ('Dependent on either isoform'; purple), reversed specifically by *Hif1a* deletion ('Dependent on HIF1A alone'; brown), reversed specifically by *Epas1* deletion ('Dependent on HIF2A alone'; green), or not reversed by any individual or combined HIFα deletion ('Dependent on neither isoform'; grey).

#### Supplementary Fig. 6

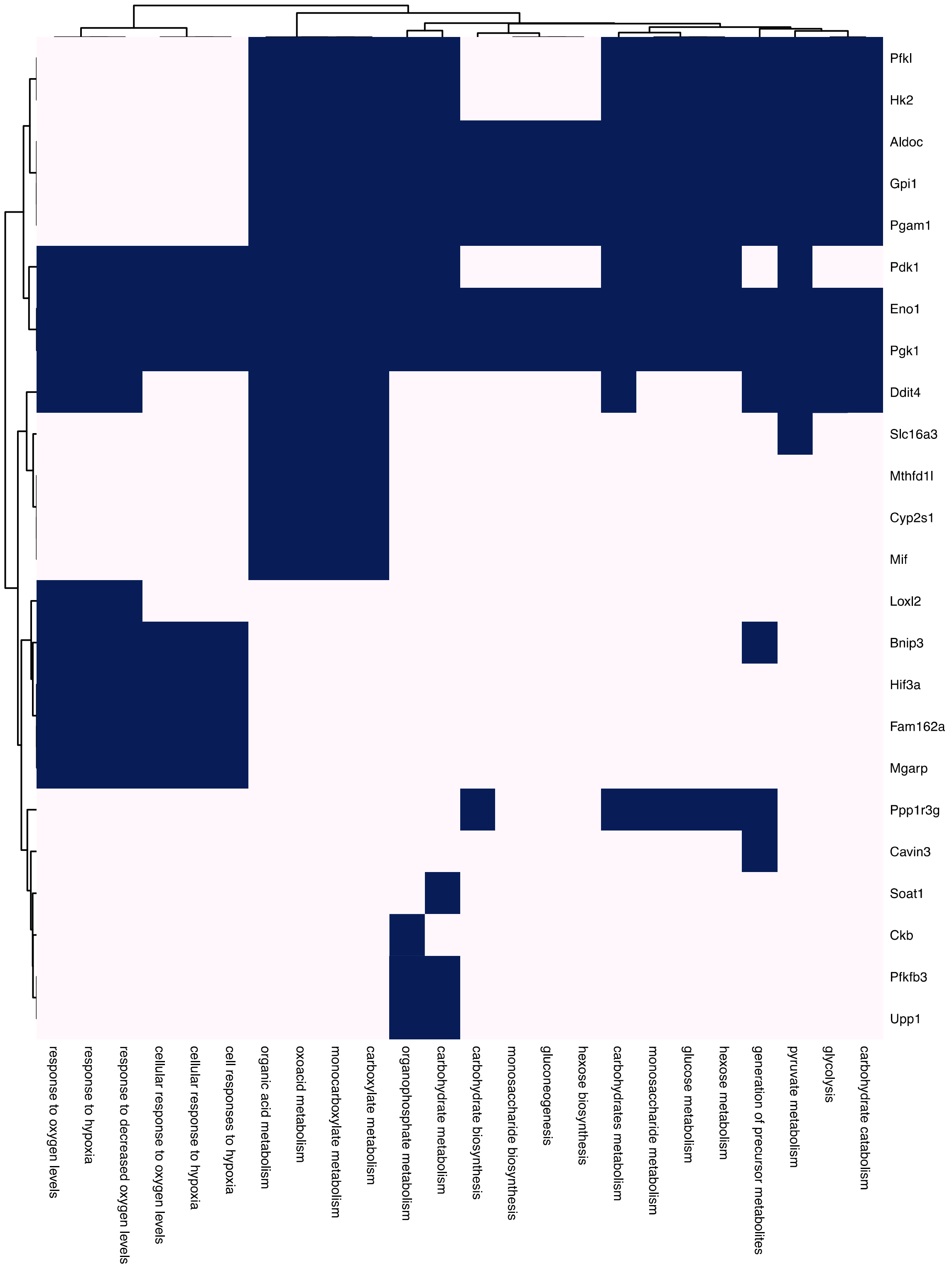


**Supplementary Fig. 6**: **Individual HIF1A-specific upregulated genes driving over-representation in the GO terms.** Binary heatmap depicting whether individual HIF1A-specific upregulated genes are or are not members of the indicated GO terms. Blue colored tiles represent membership of the gene in the GO term. GO terms and genes are ordered by hierarchical clustering based on the overlapping memberships of GO terms.

#### Supplementary Fig. 7

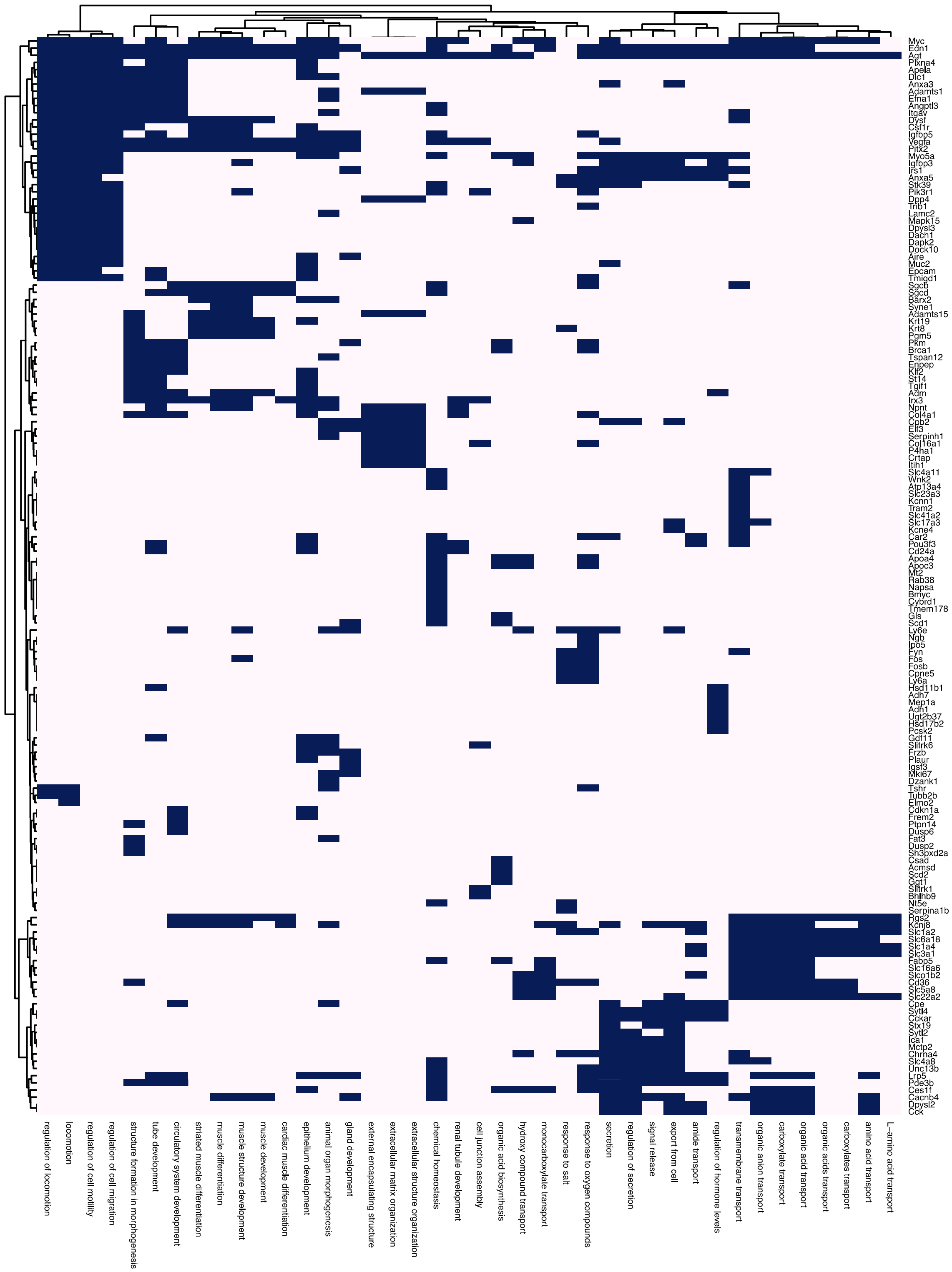

**Supplementary Fig. 7**: **Individual HIF2A-specific upregulated genes driving over-representation in the GO terms.** Binary heatmap depicting whether individual HIF2A-specific upregulated genes are or are not members of the indicated GO terms. Blue colored tiles represent membership of the gene in the GO term. GO terms and genes are ordered by hierarchical clustering based on the overlapping memberships of GO terms.

### Supplementary Fig. 8

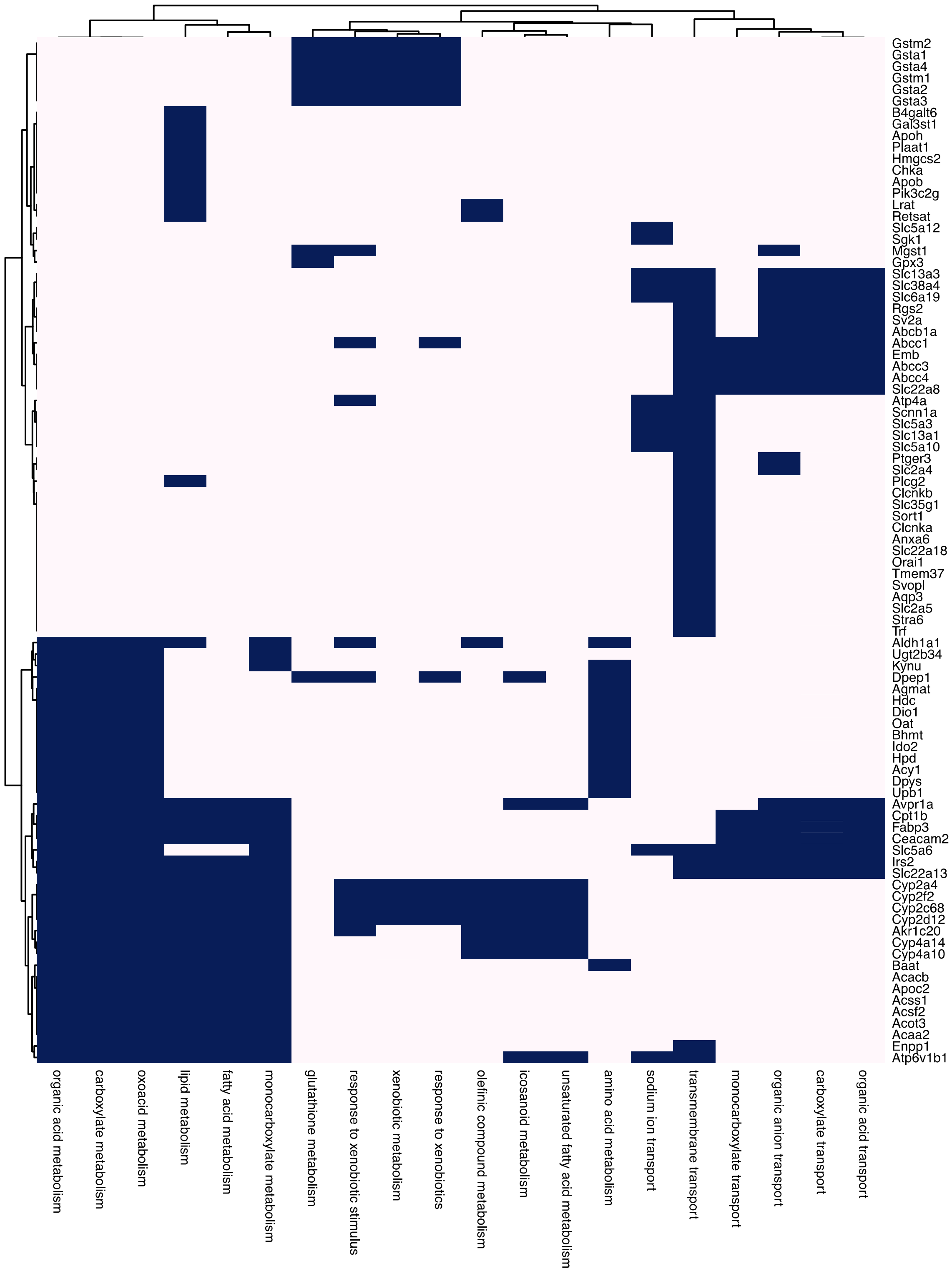


**Supplementary Fig. 8: Individual HIF2A-specific downregulated genes driving over-representation in the GO terms.** Binary heatmap depicting whether individual HIF2A-specific downregulated genes are or are not members of the indicated GO terms. Blue colored tiles represent membership of the gene in the GO term. GO terms and genes are ordered by hierarchical clustering based on the overlapping memberships of GO terms.
